# Effects and Feasibility of Hyperthermic Baths for Patients with Depressive Disorder: A Randomized Controlled Clinical Pilot Trial

**DOI:** 10.1101/409276

**Authors:** Johannes Naumann, Catharina Sadaghiani, Iris Kruza, Luisa Denkel, Gunver Kienle, Roman Huber

## Abstract

**Background:** Evaluation of efficacy, safety and feasibility of hyperthermic baths (HTB; head-out-of-water-immersion in 40°C), twice a week, compared to a physical exercise program (PEP; moderate intensity aerobic exercises) in moderate to severe depression.

**Method:** Single-site, open-label randomized controlled 8-week parallel-group pilot study at an university outpatient clinic as part of usual depression care. Medically stable outpatients with depressive disorder (ICD-10: F32/F33) as determined by the 17-item Hamilton Depression Rating Scale (HAM-D) score ≥18 and a score ≥2 on item 1 (Depressed Mood) were randomly assigned to receive either two sessions of HTB or PEP per week (40-45 min) provided by two trained doctoral students. An independent biometric center used computer-generated tables to allocate treatments. Primary outcome measure was the change in HAM-D total score from baseline (T0) to the 2-week time point (T1). Linear regression analyses, adjusted for baseline values, were performed to estimate intervention effects on an intention-to-treat (ITT) principle.

**Findings:** 45 patients (HTB *n* = 22; PEP *n* = 23) were randomized and analyzed according to ITT (mean age = 48.4 years, SD = 11.3, mean HAM-D score = 21.7, SD = 3.2). Baseline-adjusted mean difference was 4.3 points in the HAM-D score in favor of HTB (*p*<0.001). This improvement was achieved after two weeks. Compliance with the intervention and follow-up was far better in the HTB group (2 vs 13 dropouts). There were no treatment-related serious adverse events. Main limitation: the number of dropouts in the PEP group (13 of 23) was far higher than in other trials investigating exercise in depression (18.1 % dropouts).

**Conclusions:** HTB seems to be a fast-acting, safe and easy accessible method leading to clinically relevant improvement in depressive disorder after two weeks; it is also suitable for persons who have problems performing exercise training.

**Trial registration:** German Clinical Trials Register (DRKS) with the registration number <u>DRKS00011013</u> (registration date 2016-09-19) before onset of the study.

## Introduction

Depression contributes to significant economic burden and is associated with comorbid diseases (i.e. cardiovascular disease), and impaired health-related quality of life and functioning (1–5). Despite advances in the treatment of depression, one-third of depressed patients fail to respond to conventional antidepressant medication (6). Moreover, current medications cause significant side effects in the central-nervous system and commonly used antidepressants have a delayed onset of action, further highlighting the need for faster acting, easy available and more effective treatments with fewer side effects (7–9).

Fever, respectively hyperthermia, has been used as a medical treatment since ancient times, and beneficial effects of fever on mental illness were already described in antiquity (10). Based on recent findings, there is growing scientific evidence that hyperthermic baths (HTB) and whole body hyperthermia (WBH) might be efficacious for treatment of depressive disorders (11–14). The results of a non-controlled HTB study with 20 depressive patients showed an improvement in the 21-item Hamilton Depression Rating Scale (15) after five baths (11). HTB (especially before bedtime) improved sleep in healthy subjects (16–19), insomniac people (20,21) and elderly patients with vascular dementia (22). In a further non-controlled study using a radiant system to induce WBH, a single session showed a significant reduction in the Centers for Epidemiologic Studies Depression Scale (CES-D) (23) in 16 depressive patients (14). In a randomized, sham-controlled study from the same group this favorable result of a single WBH session could be corroborated (12). Our results of a randomized placebo-controlled pilot study using HTB compared with a sham light treatment were also promising; after four interventions, the intention-to-treat analysis showed a significant (*p* = 0.037) difference in the 17-item Hamilton Depression Rating Scale (HAM-D) total score of 3.14 points in favor of the HTB group (13). These results suggest that HTB and other forms of WBH can have antidepressant effects. They seem to be mediated through changes in circadian rhythm, temperature physiology and sleep, which are disturbed in depressive patients (24–26). Body core temperature in depressed patients is elevated during the night, while sleep quality is best when the core body temperature decreases; thus, change of body temperature might improve sleep quality. Findings from experimental studies show that manipulation of core and skin temperatures can improve or disrupt sleep, and it is well-known that sleep disruption negatively influences quality of life (25,27–29). Thus, sleep disturbances are an important mechanism contributing to depression (29,30). In addition, a novel hypothesis describes that an evolutionarily ancient thermoafferent pathway, signaling from serotonergic sensory cells in the skin (Merkel) and other epithelial linings to serotonergic neurons and depression-related circuits in the brain, is dysfunctional in depressive patients, might explain the antidepressant effects of HTB by restoring its function (7).

After we found HTB to be superior to a sham light therapy (13), our primary objective was to compare the efficacy of HTB with a non-pharmacological physical exercise program (PEP), a proven effective standard intervention, according to guidelines for depression (31,32). Secondary objectives were to evaluate safety (incidence of treatment discontinuation and adverse events (AE)) and feasibility of HTB applied in a home-setting.

## Method

### Ethics statement

Ethical approval was obtained from the local Ethics Committee (Ethics-Commission Medical Center University of Freiburg; 186/16; 2016-07-04).

### Study Design and Procedures

Eight-week single-site, parallel-group, open-label randomized controlled pilot trial of HTB vs PEP for patients with a diagnosis of depression according to ICD-10 (F32/F33) of at least four weeks duration. Patients were recruited from the Medical Center - University of Freiburg. The study was registered in the German Clinical Trials Register (DRKS) with the registration number <u>DRKS00011013</u> (registration date 2016-09-19) before onset of the study. The study was conducted in accordance with the Declaration of Helsinki and local laws and regulations. All the participants filled in a written informed consent form before entering the study.

### Participants

#### Inclusion criteria

Eligible participants were to meet the following criteria: (1) Medically stable outpatients with a diagnosis of depressive disorder (ICD-10: F32/F33; criteria for single or recurrent depression without psychotic features) confirmed by a physician or psychotherapist; (2) men and women between18 and 65 years of age; (3) a moderate level of depressive symptoms assessed with the 17-item Hamilton Depression Rating Scale (HAM-D) total score ≥18 and a score ≥2 on item 1 (Depressed Mood) at screening and at baseline; (4) on a consistent antidepressant regimen or off antidepressant therapy for at least four weeks prior to baseline; (5) no changes in antidepressant treatment to be expected during the study.

#### Exclusion criteria

Exclusion criteria included the presence of severe concomitant disease, epileptic disorders, organic psychotic disorders, schizophrenia, hallucinations, bipolar disorders, dissociative personality disorder, suicidal thoughts, abuse of alcohol or other drugs within the last six months, use of ß-blockers or corticosteroids, open wounds, heat urticaria, pregnancy, lactation, aversion to hot baths and participation in clinical trials in the eight weeks preceding the study. We excluded patients over age 65 because HTB causes circulatory stress.

#### Baseline assessment

Severity of depressive symptoms was assessed with the interview-based HAM-D. Patients’ subjective perception of depression severity was assessed with the self-report Beck Depression Inventory II (BDI-II) (33). Sleep quality was assessed with the self-report Pittsburgh Sleep Quality Index (PSQI) (34). Demographic and other clinical data were also collected as part of the baseline assessment, such as gender, age, duration of depression, use of antidepressant drugs, history of depression treatment including past pharmacotherapy and hospital treatment, psychotherapy (yes/no); current number of baths and sports activity in everyday life.

#### Randomization and blinding

All eligible patients who consented to participate were allocated to HTB or PEP after baseline assessment in a 1:1 ratio by simple randomization without blocking or stratification. Randomization codes were computer-generated by an independent biometric center. Allocation was performed with opaque sealed envelopes that were randomly chosen by the participants. Enrolment (RH) was done directly after informed consent was obtained from the patients. Both therapies could not be blinded. Outcome assessment (IK, LD) was unblinded. Data management and analyses were performed blinded to treatment allocation.

#### Intervention phase (Eight weeks)

Patients were randomly assigned to receive either HTB or PEP for eight weeks with two interventions per week (Table S1). Patients were told that two promising treatments were being compared in the study.

#### Hyperthermic baths (HTB)

HTB were applied as head-out-of-water-immersion in a 40 °C pool at a spa center near Freiburg, Germany. All the baths were taken in the afternoon (14:00–18:00). Five patients could sit in the pool at a time. The baths were taken until the patients noticed discomfort, the target being 20 min. Directly after the bath, the patients were accompanied to a nearby resting room, where they lay down on a resting lounger wrapped in warm blankets with two conventional 0.7 l hot water bottles (abdomen, thighs) filled with boiling water (ca. 70 °C) for at least another 20 min to keep the body temperature elevated. After 20 min in a pool with a water temperature of 40 °C, a raise in core body temperature of 1.7 °C is to be expected (35).

After four sessions under supervision (IK, LD), the remaining 12 sessions were performed by the patients themselves in the home-setting or at the spa center. At home, the baths were taken in a similar way, each bath with a duration and temperature that caused no discomfort or orthostatic reaction as tested with the first four baths. The supervisors documented the first four baths; the remaining 12 baths were documented by the patients themselves in prepared diaries.

#### Physical exercise program (PEP)

Patients in the PEP group took part in a structured physical exercise program of moderate intensity, consisting of warming-up, walking, jogging, stretching and strengthening elements for about 45-50 min, conducted outside in small groups of five patients each (for details see Table S2). The first four sessions took place under supervision (IK, LD; trained in exercise therapy). The following 12 sessions were performed by the patients themselves in groups or alone. Training duration and the exercise components completed were controlled and documented in prepared diaries.

#### Core body temperature

Core body temperature was measured with an infrared-ear-thermometer (Thermoscan^®^, Type: 6021, Braun GmbH). In the HTB group, the core body temperature was measured directly before and after the bath and after resting; in the PEP group directly before and after the exercise sessions.

### Outcome Measures

#### Primary and secondary outcomes

Unblinded assessments (IK, LD received prior training and were supervised by RH) were performed at the following three time points (for details see Table S1): before start of HTB treatment (T0), immediately on completion of the two-week treatment interval (T1; after four treatments), according to results that effects are supposed to appear early (24) and at the end of treatment (T2; after 16 treatments).

The primary outcome was determined to be the change in HAM-D total score at T1 relative to T0. Further prespecified clinical secondary outcomes were: Severity and change in scores of subjective depression symptoms as measured by the BDI-II; sleep quality as assessed with the self-report PSQI after two and eight weeks.

#### Global judgment of efficacy and tolerability

After four treatments (T1) and after end of treatment (T2), patients were asked to rate the efficacy and tolerability of the intervention on a 5-point scale (1 = very good; 2 = good; 3 = moderate; 4 = absent; 5 = worsening).

#### Safety and feasibility outcomes

To estimate the safety of HTB, the incidence of treatment discontinuation and the occurrence of AE were evaluated. To answer the feasibility question, information regarding compliance and experiences of the patients were evaluated. In order to improve adherence to intervention procedures contact was made by phone after four and six weeks, respectively.

#### Adverse events

All AE reported spontaneously by patients or observed by the assistants were recorded before and after each treatment. If a serious adverse event (SAE) occurred, the principal investigator took all the necessary and appropriate measures to ensure the safety of the patient.

### Data Analysis

#### Sample size estimation

No data on effect size of HTB in comparison to PEP were available; however, from the results of our previous study, we expected a moderate to large effect size for HTB. An effect size of *d* = 0.91 and 1-β = 0.80 power was chosen for the study, which required 40 patients (n = 20 per arm) to reach significance with α = 0.05. This estimation was based on the primary outcome of depression symptoms as measured by the HAM-D total score at two weeks of intervention (36).

#### Statistical analyses

Treatment comparison of the primary endpoint (HAM-D change from baseline to T1) was performed within a generalized linear regression model adjusted for baseline values. BDI-II and PSQI were analyzed in the same way. Analyses were done on the intention-to-treat (ITT) population, defined as all allocated patients, applying the last-observation-carried-forward (LOCF) approach to impute missing data. Baseline characteristics were compared using 2-sided *t* tests for continuous data and 𝒳 ^2^ statistics. The per-protocol (PP) population was defined as all patients who had a complete dataset for the primary outcome and had participated in at least 75 % of the treatments, meaning at least 3 of 4 treatments for T1, and at least 12 of 16 treatments for T2. We report *p*-values with the significance level set at *p*<0.05. The study followed a protocol (Protocol S1-S3) and reporting followed the CONSORT (Consolidated Standards of Reporting Trials) Statement extension for nonpharmacological treatments (37). Prespecified secondary analyses were not adjusted for multiple comparisons and should therefore be regarded as descriptive and exploratory. Statistical analyses were performed using SPSS^®^, Version 24, for WindowsTM.

#### Data collection and monitoring

A data manager (CS), blind to treatment allocation, reviewed and evaluated data, in order to detect errors during data collection, and conducted a quality review of the database, with double data entry by two independent persons for 20 % of the values.

## Results

### Enrolment and Sample Description

The HTB study began recruiting patients in September 2016 and closed recruitment in January 2017. 69 adults agreed to participate and were assessed for eligibility. Of that number, 45 underwent randomization (Figure 1).

**Figure 1.**
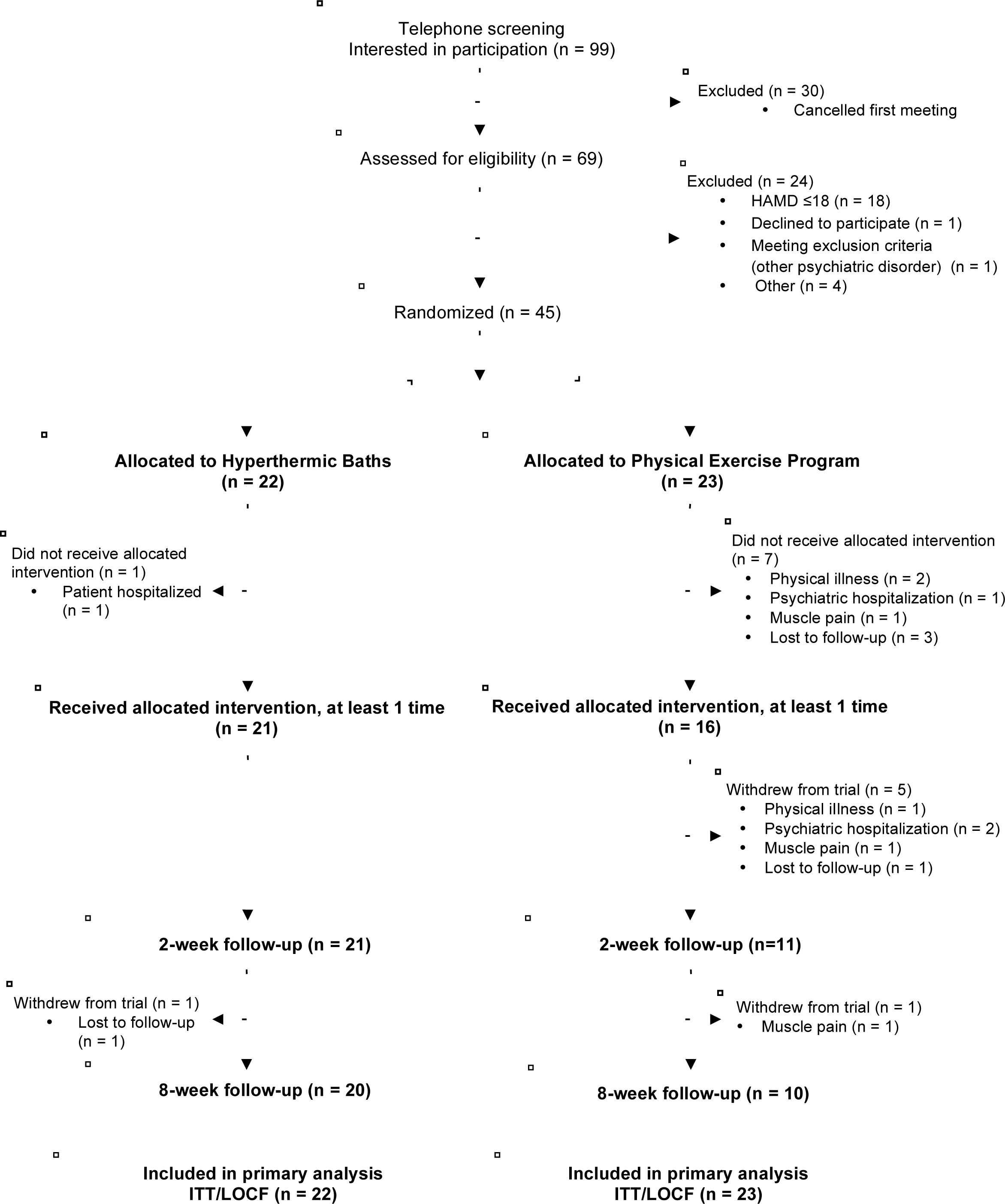
CONSORT flow diagram of study participants.

Randomization was balanced with respect to demographic and clinical characteristics with more females (*p* = 0.047), a higher BMI (*p* = 0.135) and longer duration of depressive disorder in the PEP-group (*p* = 0.035), (Table 1). The mean depression severity score based on the HAM-D was 21 (HTB), respectively 22 (PEP), consistent with moderate severity depression (38). Self-reported severity and symptoms of depression assessed by the BDI-II was 29 (HTB), respectively 31 (PEP), consistent with the lowest possible score for severe depression. Most patients had depression for several years; the shortest duration of depression was two months, the longest more than 20 years. Sleep quality, as measured by the PSQI total score (ranging from 0-21; score ≤5 associated with good sleep quality; score >5 associated with poor sleep quality), was poor with a total score of 10 (HTB), respectively 11 (PEP). Prescription rates of antidepressants were high at baseline (HTB, 68 %; PEP, 48 %), as well as for psychotherapy (HTB, 71 %; PEP, 83 %).

Primary outcome data (HAM-D total score after two weeks) were available for 32 patients, representing a loss to follow-up of 29 % (5 % in the HTB group and 52% in the PEP group). Eight patients (HTB n = 1; PEP n = 7) discontinued treatment without receiving their allocated intervention. A further five patients (PEP group) withdrew from trial before T1; another two after T1 (HTB n = 1; PEP n = 1).

**Table 1.** Demographic and clinical characteristics at baseline.

|  | **Overall**<br>**(N = 45)** | **Hyperthermic**<br>**Baths**<br>**(N = 22)** | **Physical Exercise**<br>**Program**<br>**(N = 23)** |
| --- | --- | --- | --- |
| Gender: Female, N (%) | 33 (73%) | 13 (59%) | 20 (87%) |
| Age (years) | 48.4 (11.3) | 45.6 (12.3) | 51.0 (9.8) |
| BMI (kg/m<sup>2</sup>) | 25 (5) | 24 (4.2)<sup>\$\$</sup> | 26 (5.6)<sup>\$</sup> |
| Median (range) | 24.5 (16-35) | 23.5 (17-33) | 26 (16-35) |
| Duration of depression, years | 6 (7) | 4 (6) | 8 (7,5) |
| Psychiatric hospital stays (n)<br>(last 2 years) | 21 (48%) | 11 (52%)<sup>\$</sup> | 10 (43%) |
| Use of psychopharmaca, n (%) | 26 (58%) | 15 (68%) | 11 (48%) |
| Psychotherapy | 34 (77%) | 15 (71%)<sup>\$</sup> | 19 (83%) |
| HAM-D total score | 21.7 (3.2) | 21.2 (3.3) | 22.2 (3.1) |
| Median (range) | 21 (18-29) | 21 (18-29) | 22 (18-29) |
| BDI-II | 30.0 (7.1) | 29.0 (5.8) | 31.1 (8.1) |
| Median (range) | 31 (13-50) | 30.5 (19-38) | 31 (13-50) |
| PSQI | 10.8 (3.7) | 10.2 (3.8) | 11.4 (3.7) |
| Median (range) | 10 (4-18) | 10 (4-16) | 11 (5-18) |

BMI, Body Mass Index; HAM-D, 17-item Hamilton Depression Rating Scale; BDI-II, Beck Depression Inventar II; PSQI, Pittsburgh Sleep Quality Index.
<sup>\$</sup>missing *n* = 1; <sup>\$\$</sup>missing *n* = 2;
Where not otherwise indicated, data are shown as mean and standard deviation (SD).

#### Treatment effect on core body temperature

A bath taken for up to 30 min at a water temperature of 40 °C resulted in a rise of core body temperature from 36.7 °C before the bath to 38.6 °C directly after the bath (mean change 1.96, standard deviation [SD] 0.5, and in it being maintained at 37.4 °C (mean change 0.7, SD = 0.5) after rest. In the PEP group, there was no difference in core body temperature before or after physical exercise (36.48 °C, SD = 0.46, respectively 36.50 °C, SD = 0.52).

### Primary Outcome

The ITT analysis showed a significant adjusted mean difference between the groups of 4.3 points (95% CI 2.16 to 6.42) in the HAM-D score after two weeks in favor of HTB (p<0.001, Table 2). In PP analyses (≥75% adherence), this difference was only found as a trend (*p* = 0.068, Table 3).

**Table 2.** Between-group differences in the intent-to-treat last observation carried forward sample.

| Outcome Measure | HTB Group<br>(N = 22) | PEP Group<br>(N = 23) | HTB vs PEP<br>Adjusted mean difference<sup>§</sup><br>(95% CI) | p<sup>§</sup> |
| --- | --- | --- | --- | --- |
| HAM-D total score |  |  |  |  |
| Baseline | 21.2 (3.3) | 22.2 (3.1) |  |  |
| T1: 2 weeks<sup>1</sup> | 15.8 (4.4) | 20.8 (4.2) | 4.3 (2.16 to 6.42) | **<0.001** |
| T2: 8 weeks<sup>2</sup> | 15.3 (5.8) | 19.0 (6.1) | 2.9 (-0.40 to 6.14) | 0.084 |
| BDI-II |  |  |  |  |
| Baseline | 29.0 (5.8) | 31.1 (8.1) |  |  |
| T1: 2 weeks<sup>2</sup> | 20.3 (8) | 29.1 (8.8) | 7.5 (2.99 to 11.93) | **0.002** |
| T2: 8 weeks<sup>2</sup> | 20.5 (10.8) | 24.8 (11.8) | 3.1 (-3.36 to 9.53) | 0.340 |
| PSQI |  |  |  |  |
| Baseline | 10.2 (3.8) | 11.4 (3.7) |  |  |
| T1: 2 weeks<sup>2</sup> | 8.7 (3.7) | 11.5 (4.1) | 2.0 (0.14 to 3.92) | **0.036** |
| T2: 8 weeks<sup>2</sup> | 8.3 (3.2) | 10.7 (4.1) | 1.8 (-0.13 to 3.64) | 0.067 |

Abbreviations: HAM-D, 17-Item Hamilton Depression Rating Scale; BDI-II, Becks Depression Inventar II; PSQI, Pittsburgh Sleep Quality Index.
Data are shown as mean and standard deviation (SD).
<sup>1</sup>Primary outcome.
<sup>2</sup>Prespecified secondary outcome.
<sup>§</sup>Linear regression analyses adjusted for baseline scores.

**Table 3.** Between-group differences in the per-protocol completer sample.

| Outcome Measure | HTB Group | PEP Group | HTB vs PEP<br>Adjusted mean difference<sup>§</sup><br>(95% CI) |  | p<sup>§</sup> |
| --- | --- | --- | --- | --- | --- |
| HAM-D total score |  |  |  |  |  |
| Baseline | 21.0 (3.2) | 21.4 (3.0) |  |  |  |
| T1: 2 weeks* | 14.9 (3.6) | 17.9 (4.2) | 2.7 | (-0.22 to 5.69) | 0.068 |
| T2: 8 weeks** | 14.1 (5.2) | 12.9 (5.1) | -0.9 | (-5.66 to 3.97) | 0.72 |
| BDI-II |  |  |  |  |  |
| Baseline | 28.5 (6.1) | 32.0 (4.1) |  |  |  |
| T1: 2 weeks* | 19.2 (7.8) | 24.2 (6.9) | 3.0 | (-3.09 to 9.08) | 0.32 |
| T2: 8 weeks** | 18.0 (10.7) | 13.0 (6.5) | -7.7 | (-16.19 to 0.76) | 0.07 |
| PSQI |  |  |  |  |  |
| Baseline | 9.9 (3.9) | 10.6 (3.0) |  |  |  |
| T1: 2 weeks* | 8.3 (3.8) | 9.6 (3.9) | 0.9 | (-1.83 to 3.57) | 0.51 |
| T2: 8 weeks** | 8.1 (3.6) | 8.0 (3.9) | -1.2 | (-3.74 to 1.39) | 0.35 |

\* HTB *n* = 19; PEP *n* = 9; \*\* HTB *n* = 15; PEP *n* = 7
Abbreviations: HAM-D, 17-Item Hamilton Depression Rating Scale; BDI-II, Becks Depression Inventar II; PSQI, Pittsburgh Sleep Quality Index.
Data are shown as mean and standard deviation (SD).
<sup>§</sup>Linear regression analyses adjusted for baseline scores.

### Descriptive Secondary Analyses

Looking at the mean differences compared to baseline, the HAM-D results show a stable improvement in the HTB group of 5.4 points after two weeks and of 5.9 points after eight weeks, whereas in the PEP group we see an improvement of 1.4 points after two weeks and of 3.2 points after eight weeks (Figure 2A).

A similar pattern can be seen in the results of the BDI-II, with a mean difference compared to baseline in the HTB group of 8.7 points after two weeks and of 8.5 points after eight weeks. The PEP group showed an improvement of 2.0 points after two weeks and of 6.3 points after eight weeks (Figure 2B). Results of the PSQI show a mean difference compared to baseline of 1.5 points after two weeks and of 1.9 points after eight weeks in the HTB group, and a deterioration of 0.1 points after two weeks and an improvement of 0.7 points after eight weeks in the PEP group (Figure 2C).

**FIGURE 2.**
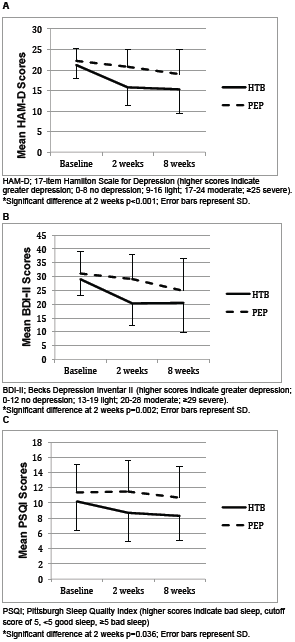
Modeled change scores by week.^*^

Subgroup-analyses according to HAM-D score quartiles (median) revealed the greatest treatment effect in quartiles 3 and 4. We found a difference of 7.9 (SD = 3.7) for HAM-D ≥21 at baseline in the HTB group (*n* = 10) and of 3.4 (SD = 2.1) in the PEP group (*n* = 5) and a difference of 11.5 (SD = 2.7) for HAM-D ≥24 at baseline in the HTB group (*n* = 4) and of 3.0 (SD = 0) in the PEP group (*n* = 2).

#### Global judgment of efficacy and tolerability

The global judgment by the patients showed no significant differences between the groups, with good to moderate efficacy (HTB 2.6 (SD = 1.2); PEP 2.4 (SD = 1.4); *p*_2-tailed_ = 0.76) and tolerability (HTB 2.1 (SD = 1.0); PEP 1.6 (SD = 0.7); *p*_2-tailed_ = 0.16) after four interventions and good to moderate efficacy (HTB 2.1 (SD = 1.0); PEP 2.3 (SD = 1.1); *p*_2-tailed_ = 0.53) and tolerability (HTB 1.8 (SD = 0.8); PEP 1.7 (SD = 0.7); *p*_2-tailed_ = 0.66) at the end of treatment after 16 interventions.

#### Adverse events

Adverse events possibly related to the therapy were reported by 25 patients, of which 18/21 were assigned to the HTB group and 7/11 to the PEP group (Table S3). No SAE were reported by either group. There was no significant difference between the groups (*p*_2-tailed_ = 0.197). Typical AE in the HTB group were discomfort during the baths such as dizziness, fatigue, palpitations and thus mainly attributable to the cardiovascular system and indicating that the HTB were applied at a therapeutic limit. Additionally, patients reported transient effects as minor headache, itching, sweating, tightness, hunger worsening of depression and irritability. Typical AE in the exercise group were muscle soreness, pain in joints, the lower back or tendons; one patient reported cough and dizziness caused by breathing in cold air during exercise.

#### Dropouts and compliance

Adverse events that were not related to the treatment but which led to dropout included one patient in the HTB group who was hospitalized before the start of the treatment and six in the PEP group, three of which withdrew because of physical illness and three because of psychiatric hospitalization; three of these before the start of the treatment and the other three during the first two weeks of treatment.

Treatment related AE resulted in three dropouts in the PEP group (muscle pain); there were no treatment-related dropouts in the HTB group. Compliance was good with a medium number of treatments of 13.3 (95% CI 11.6 to 14.9) in the HTB group and of 12.6 (95% CI 9.1 to 16.1) in the PEP group.

#### Feasibility of HTB in the home-setting

After four sessions under supervision, the remaining 12 sessions were performed without supervision, either in the home-setting or at the spa center as before. The results show that HTB can be performed without supervision. However, only seven patients attempted to take the baths at home, and their main complaint was that it was difficult to reach the target water temperature of 40 °C; thus, they returned to the spa center.

## Discussion

To our knowledge, this is the first randomized controlled parallel group study to assess the efficacy of HTB compared with PEP in adults with depressive disorder. The main finding from this preliminary study is that HTB significantly reduced the HAM-D score compared with PEP. It should be emphasized that the onset of treatment response occurred within the first two weeks. Commonly used antidepressants, the standard treatment for depression, have a delayed onset of action. It requires weeks of treatment before the core symptoms of depression are ameliorated (9). In SSRIs (selective serotonin reuptake inhibitors), i.e. fluoxetine, this discrepancy between antidepressant-induced acute neurochemical effects and clinical effectiveness adds up to four weeks in 75% of responders (8). Consequently, HTB could offer a concomitant treatment option to close this gap. Exercise as a treatment for depression took six to eight weeks to show an effect (Figure 2). The between-group difference after two weeks, adjusted for baseline values, was 4.3 HAM-D points (*p*<0.001), which is a clinically relevant difference. As established by the National Institute for Clinical Excellence (NICE) the threshold for clinical relevance is a difference of 3 points on the HAM-D (39). As in pharmacological studies, the magnitude of the difference in HAM-D scores between the HTB and the PEP group increased with increasing baseline depression severity. These results are supported by the outcomes of the BDI-II, which also showed a significant and clinically relevant improvement with an adjusted mean difference of 7.5 points (*p* = 0.002), representing a decrease of 30% between the groups in favor of the HTB group after two weeks; however, after eight weeks, the difference was no longer significant (*p* = 0.34). According to the NICE-guidelines a difference of 3 points in the BDI-II, respectively a decrease of 17.5 % compared to baseline is regarded as clinically relevant (39). Whether the improvement in sleep in the HTB group after two weeks as seen in the PSQI is the cause or the consequence of the improvement in depression cannot be answered with this trial.

HAM-D total score was also lower at eight weeks in the HTB group, but high attrition rates reduce confidence in this result. The PP analysis showed a trend towards better results in the PEP group at eight weeks. This however, may be due to a selection bias as patients with severe depression were hospitalized. The patients’ overall global judgment of efficacy showed no difference between the groups. This is similar to the results from Button et al. (39), respectively the TREAD trial (40). The global rating of change was “better” only with a corresponding mean change of −18.9 in the BDI-II, whereas a mean change of −6.4 was still regarded as “no” improvement.

There were fewer females in the HTB group (59 %) than in the PEP group (87 %); however, in a recent review gender did not modify the antidepressant effect of exercise (41). In members of the PEP group, symptoms of depression had lingered for a longer period (eight years) than in those of the HBT group (four years). Because longer duration of depression negatively affects treatment outcome the difference may have influenced the results (42).

Safety concerns existing prior to the study, especially regarding orthostatic dysregulation after HTB in the unsupervised part of the study, were not confirmed. Patients in both groups were able to adapt the intensity or duration of the therapy. We found a high rate of minor AE in both groups, but there were no SAE. In conclusion, HTB performed at a spa center can be regarded as safe and feasible without external supervision, provided that the patients are informed concerning the potential risks, especially regarding orthostatic dysregulation. Whether HTB is feasible in the home-setting cannot be answered conclusively, because only a few patients bathed there. Although not under investigation, the high attrition rate in the PEP group indicates that physical exercises are not suitable for everyone. Our finding of significant efficacy of the HTB intervention confirms the results of previous studies (11–13,24).

### Strengths and limitations

The strengths of our study are the randomized, controlled design, the use of standardized baths, the good control of body temperature and the use of established patient as well as investigator-related questionnaires. Several limitations should be discussed. First, the number of dropouts in the PEP group (13 of 23) was far higher than in other trials investigating exercise in depression (18.1 % dropouts) or even in major depression (17.2 %), but with higher baseline depressive symptoms predicting greater dropout (43). This might have been due to the fact that dropout rates are likely to be higher in the outpatient than in the inpatient setting (43). On the other hand, it indicates that this study population was difficult to motivate to active treatment and that HTB may be an even better accepted form of treatment than exercise. Second, because of the small sample size, the study has limited power to detect clinically significant differences between the treatment conditions, especially in subgroup analyses. Third, the absence of blinding of treatment conditions, which is inherent and inevitable; due to unblinded outcome assessment a risk for performance bias exists. Fourth, it is a well-known fact that the HAM-D total score has pitfalls; however, for better comparability with other studies, we did not use the GRID-HAM-D, e.g., with better reliability and validity (44,45). Fifth, the LOCF method for missing values may have led to an overestimation of the treatment effect of HTB.

### Generalizability

Although external validity may be restricted due to the population selected to participate in clinical studies, the population studied here can be regarded as representative of routine clinical practice, including patients with and without antidepressant medication (46). Contraindications to HTB are still not well defined. Severe concomitant diseases, i.e. cardiovascular, or orthostatic dysregulation should be omitted, especially in the elderly.

## Conclusion

HTB seems to be a fast-acting and safe method leading to clinically relevant improvement in depressive symptoms after just two weeks. Patients can apply the method at their own responsibility and it can also be practiced by patients with problems performing exercise training.

Replication of the results in a large, confirmative trial would clearly have important implications for public health in that the emotional and financial tolls of moderate to severe depression could be mitigated with a low-cost and easily accessible intervention with high acceptance by patients.

## Acknowledgments

The authors are grateful to all the participants of the study. We thank Tania Lüty for assistance in preparing the manuscript.

The article processing charge was funded by the German Research Foundation (DFG) and the University of Freiburg in the funding programme Open Access Publishing.

Margarete Müller-Bull Stiftung, Gerokstr. 1, 70188 Stuttgart, Germany http://www.mmb-stiftung.de/deutsch/kontakt/impressum.html (Uni-Zentrum Naturheilkunde)

The funders had no role in study design, data collection and analysis, decision to publish, or preparation of the manuscript.

## Supporting information

Supplement (PDF):

- TABLE S1. Diagram of the study protocol

- TABLE S2. Outline of Physical Exercise Program (PEP)

- TABLE S3. Summary of adverse events.

Protocols (PDF):

- S1: Study protocol (German)

- S2: Study protocol (English, key methodological section)

- S3: DRKS registration

Text S1 (DOC): CONSORT extension for pilot and feasibility trials checklist

## Abbreviations

BDI-II: Beck Depression Inventar II
CES-D: Centers for Epidemiologic Studies Depression Scale
CI: confidence interval
DRKS: German Clinical Trial Register
HAM-D: 17-item Hamilton Scale for Depression
HTB: hyperthermic baths
ITT: intention-to-treat
LOCF: last-observation-carried-forward
PEP: Physical Exercise Program
PP: per-protocol
PSQI: Pittsburgh Sleep Quality Index
SD: standard deviation
WBH: whole body hyperthermia.

